# Characterization of tri-culture alveolar-like organoids to study fibrotic lung diseases

**DOI:** 10.64898/2026.09.04.749305

**Authors:** Tony J.F. Guo, Yidan Meng, Seowon Kim, Tak Machiri, Allyson Miscampbell, Kevin S.K. Lau, Clara Gibbons, Mackenzie R. Penton, Charlotte P.S. Lee, Isaac T.S. Li, Gillian C. Goobie, Andrew J. Halayko, Christopher J. Ryerson, Christopher Carlsten, Min Hyung Ryu, Emmanuel T. Osei

## Abstract

**Rationale:** Fibrotic interstitial lung diseases are progressive disorders characterized by lung scarring and declining respiratory function. Repeated injury and dysregulated epithelial-mesenchymal crosstalk are implicated in disease pathogenesis but remain incompletely understood. Three-dimensional (3D) organoids incorporating epithelial-mesenchymal crosstalk provide a physiologically relevant platform for investigating these mechanisms. Here, we present a 3D tri-culture alveolar organoid model and evaluate its responses to two profibrotic stimuli, TGF-β1 and bleomycin.

**Methods:** CI-huArlo, NCI-H441, and MRC-5 cells were co-cultured for 14 days to generate alveolar-like organoids. Cell marker expression was assessed by immunofluorescence (IF). To test injury response, organoids were exposed to TGF-β1 (50 ng/mL) or bleomycin (20 μg/mL) for 48 hours and characterized. Supernatants were collected to assess interleukin 8 (IL-8) and procollagen I by ELISA. qPCR assessed expression of *CDKN1A* and *COL1A1* following treatments. Bulk RNA sequencing (RNA-seq) evaluated the transcriptomic responses to fibrotic stimulus.

**Results:** Cellular marker expression was confirmed using IF for aquaporin-5 (alveolar type I cell marker) and TE-7 (fibroblast marker). Bleomycin exposure reduced viability and significantly increased IL-8 (126.3 ± 16.86 vs. 48.88 ± 4.470 pg/mL; p = 0.0002; N=6), with no change in secreted procollagen I compared to control. TGF-β1 stimulation significantly increased secreted procollagen I (149.6 ± 27.07 vs. 53.38 ± 5.672 pg/mL; p <0.0001; N=6) without affecting viability or IL-8 release. qPCR resulted in no change in *CDKN1A* expression, while *COL1A1* expression was increased with TGF-β1 treatment compared with control and bleomycin. RNA sequencing demonstrated distinct and reproducible transcriptional responses to TGF-β1 and bleomycin. TGF-β1 induced 750 significantly upregulated and 641 downregulated genes, including increased *COL1A1*, *COL4A1*, *FN1*, and *TGFB1*, with enrichment of epithelial-mesenchymal transition and TGF-β signalling programs. In contrast, bleomycin induced 953 significantly upregulated and 552 downregulated genes relative to untreated controls and was characterized by p53 signalling, DNA-damage responses, and reduced cell-cycle progression. Reference-state analysis further indicated reduced normal alveolar epithelial signatures following both treatments, with TGF-β1 producing the strongest aberrant basaloid and myofibroblast-associated signatures.

**Conclusions:** Tri-culture alveolar-like organoids exhibited stimulus-specific responses in a 3D multicellular system, supporting their use for mechanistic studies of epithelial-mesenchymal crosstalk, environmental exposures, and therapeutic responses.

## Introduction

Fibrotic interstitial lung diseases (fILDs), including idiopathic pulmonary fibrosis (IPF) and other ILD subtypes, are progressive disorders characterized by persistent scarring that disrupts alveolar architecture, impairs gas exchange, and contributes to respiratory decline and mortality [1–4]. This clinical burden, together with limitations of existing experimental models in studying human fibrotic lung biology, supports the need for complementary human relevant models [5].

The pathogenesis of fILD is thought to involve repeated alveolar epithelial injury followed by dysregulated repair [6]. Genetic susceptibility also contributes to disease development in IPF [7]. A *MUC5B* promoter variant is associated with pulmonary fibrosis, while variants affecting telomere maintenance and surfactant homeostasis are associated with additional pathways that predispose individuals to epithelial dysfunction and fibrotic remodelling [7–10]. Recurrent alveolar epithelial injury from environmental and occupational exposures, including inhaled dusts and other particulates, contributes to disease initiation and progression in genetically susceptible individuals, although specific gene-environment interactions remain incompletely defined [11]. In healthy lung tissue, the alveolar epithelium, fibroblasts, immune cells, and extracellular matrix (ECM) work together to maintain tissue structure and restore barrier integrity after injury [12]. In fibrotic disease, this repair process becomes maladaptive. Injured epithelial cells release inflammatory and profibrotic mediators that can activate fibroblasts and promote excessive ECM deposition [13]. Over time, these processes create a fibrotic niche, a dynamic microenvironment in which epithelial dysfunction, fibroblast activation, inflammatory signalling, and matrix remodelling reinforce abnormal tissue repair [13].

Epithelial-mesenchymal crosstalk is central to this process. Alveolar type 1 (AT1) and type 2 (AT2) epithelial cells are targets of injury and regulators of the fibrotic response, while fibroblasts respond to epithelial-derived signals by changing their behaviour, matrix production, and inflammatory mediator release. Profibrotic stimuli such as transforming growth factor-β1 (TGF-β1) can promote ECM production, while injury-inducing agents such as bleomycin can trigger epithelial stress, inflammation, and tissue damage responses [14,15]. Dysregulated epithelial repair in IPF is also associated with the emergence of aberrant basaloid cells, a fibrosis-associated population with basal epithelial, cellular stress, senescence, developmental, and mesenchymal-associated transcriptional programs [16–18]. These cells localized to remodelled alveolar regions and fibrotic foci, and their transcriptional signatures are associated with disease severity [19].

Two-dimensional monoculture models have been useful for individual cell types and pathways but lack three-dimensional architecture, epithelial-stromal interactions, matrix-embedded growth, and the local microenvironmental cues that influence cell behaviour *in vivo* [20]. Animal models enable investigation of fibrogenesis within an intact multicellular system, but no single model reproduces all clinical, histopathological, and temporal features of human IPF [21]. Complementary human *in vitro* models are therefore needed to study early fibrotic mechanisms, dissect gene-environment interactions, and test environmental and occupational exposures against potential therapeutic targets.

Among these models, organoids incorporate multiple cell types and extracellular matrix components within self-organizing three-dimensional structures, allowing the study of tissue-like responses that are difficult to reproduce in standard monolayer cultures [22]. Alveolar organoid models have been used to study chronic lung diseases, epithelial injury, regeneration, inflammation, and fibrosis-related pathways [23–26]. However, many existing models utilize primary or pluripotent stem cell-derived cells that require lengthy or technically complex differentiation procedures. A defined combination of established cell lines may enable shorter culture timelines and scalable comparisons while retaining epithelial-stromal interactions. In this study, we developed and characterized a tri-culture alveolar-like organoid model for studying stimulus-specific responses associated with fibrotic lung injury.

## Methods

### Cell culture

CI-huArlo immortalized human alveolar epithelial cells were obtained from InSCREENex (Braunschweig, Germany), NCI-H441 human lung papillary adenocarcinoma cells and MRC-5 fetal lung fibroblasts were obtained from ATCC (Manassas, USA). CI-huArlo cells were cultured on flasks coated with huAEC coating solution (InSCREENex) in complete PneumaCult-Ex medium (STEMCELL Technologies, Vancouver, Canada). NCI-H441 and MRC-5 cells were cultured in RPMI 1640 medium (Thermo Fisher Scientific, Waltham, MA) and high glucose DMEM (Thermo Fisher Scientific), respectively, supplemented with 10% fetal bovine serum (FBS, Thermo Fisher Scientific). All media contained 100 U/mL penicillin-streptomycin (Thermo Fisher Scientific). Cells were maintained below passage 20.

### Organoid medium generation and characterization

Terminal-airway Organoid Medium (TOM) was made using a 75/25 mixture of advanced DMEM (Thermo Fisher Scientific) and advanced DMEM/F12 (Thermo Fisher Scientific), 2% FBS, 100 U/mL penicillin-streptomycin, 20 nM of hydrocortisone (STEMCELL Technologies), and 2 mM of L-glutamine (Thermo Fisher Scientific). This formulation was adapted from previous studies [27,28]. Its suitability for the three cell lines was compared with DMEM, RPMI-1640, and PneumaCult-Ex media using a PrestoBlue viability assay (Thermo Fisher Scientific). Cells were exposed to each medium for 24 or 48 hours before incubation with 10% PrestoBlue for 2 hours. Fluorescence was measured at 560 nm excitation and 590 nm emission.

### Organoid dome generation and monitoring with light microscopy

CI-huArlo, NCI-H441, and MRC-5 cells were dissociated and counted using trypan blue exclusion staining. Each organoid dome contained 13,200 cells at a ratio of 11:7:4 of NCI-H441, CI-huArlo, and MRC-5 cells. Cells were pelleted at 250 x g for 5 minutes, resuspended in 3.5 μL of chilled TOM and 31.5 μL of GelTrex Flex (GelTrex, Thermo Fisher Scientific), and the 35 μL suspension was dispensed as domes into prewarmed 24-well suspension culture plates and polymerized for 30 minutes at 37°C. Representative images of these domes are seen in **Supp. Fig. 1**. Each well then received 500 μL of TOM containing 2.5% GelTrex. Organoids were cultured for 14 days, with medium replaced every 4 days. Domes were imaged using light microscopy every 3 days to assess for morphology on the EVOS M5000 microscope (Thermo Fisher Scientific). Organoid size and counts were quantified in ImageJ software [29] by two blinded observers.

**Figure 1.**
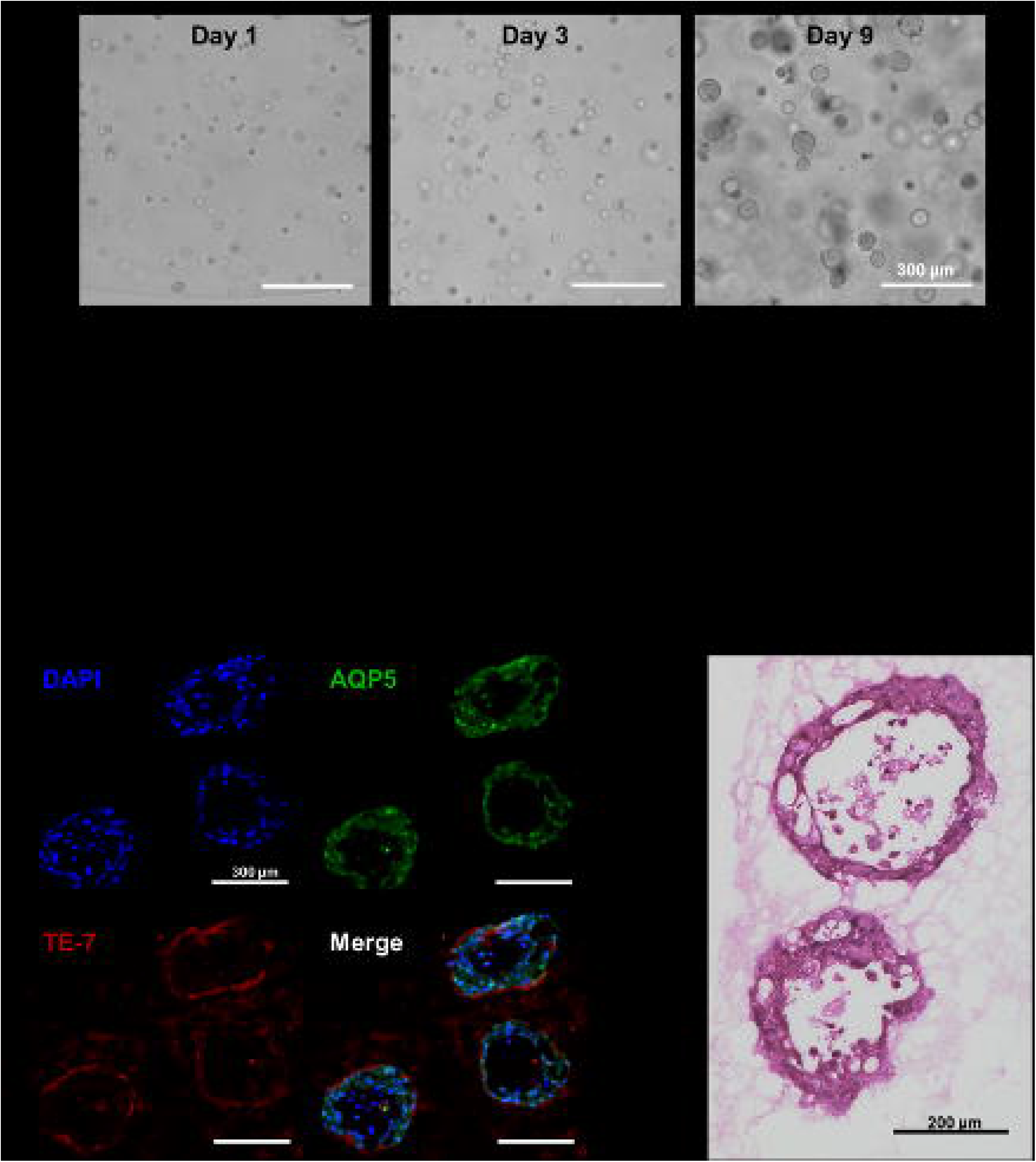
Growth and structural characterization of tri-culture alveolar-like organoids. **(A)** Representative transmitted light images of tri-culture organoids on days 1, 3, and 9 of culture showing progressive organoid formation and growth. Scale bar = 300 mm. **(B)** Quantification of organoid diameter over time with growth. **(C)** Quantification of organoid count over time with growth. **(D)** Representative immunofluorescence images of organoid sections stained for the alveolar type I cell marker, aquaporin-5 (AQP5), green, fibroblast-specific marker, TE-7, red, and nuclei, DAPI, blue, with the scale bar = 300 mm. **(E)** Representative hematoxylin and eosin (H&E) stained cross-section of cryosectioned organoids with the scale bar = 200 mm.

### Treatment of organoids with TGF-**β**1 and bleomycin sulfate

After the 14 days, organoids were recovered using organoid harvesting solution (R&D Systems, Minneapolis, USA) according to manufacturer’s instructions. The pellet obtained from each dome was passaged into two domes and resuspended in 3.5 μL of chilled TOM and 31.5 μL of GelTrex with recombinant TGF-β1 protein (Peprotech, Cranbury, USA) at 50 ng/mL or with bleomycin sulfate (Sigma-Aldrich, St. Louis, USA) at 20 μg/mL. 35 μL of the organoid-GelTrex mixture was pipetted onto a pre-warmed hydrophobic cell culture plate and incubated at 37°C for 30 minutes to solidify. To each well, 2.5% GelTrex in TOM, with TGF-β1 at 50 ng/mL or with bleomycin sulfate at 20 μg/mL in the same medium was added. The domes were returned to incubation for 48 hours at 37°C and imaged using light microscopy at 24 and 48 hours after dome formation. The supernatant was collected after 48 hours of treatment, and PrestoBlue cellular viability assay (Thermo Fisher Scientific) was performed according to manufacturer’s instructions. Organoids were subsequently collected for RNA isolation or fixed in 1% paraformaldehyde for histological analysis.

### Quantitative PCR and immunoassays

Quantitative PCR was performed using a QuantStudio 6 Pro Real-Time PCR system (Thermo Fisher Scientific). GAPDH-VIC (Assay ID: Hs99999905_m1) was used as the endogenous reference assay, with forward (5′-CAAGGTCATCCATGACAACTTTG-3′), and reverse (5′-GGCCATCCACAGTCTTCTGG-3′) primers as previously reported [30]. COL1A1-FAM (Assay ID: Hs00164004_m1) and CDKN1A-FAM (Assay ID: Hs00355782_m1) were used as target assays. Relative gene expression was calculated by normalizing target gene expression to GAPDH.

Supernatant interleukin-8 (IL-8) and pro-collagen I alpha 1 were assayed using the human IL-8/CXCL8 ELISA (DY208, R&D Systems) and the human pro-collagen I alpha 1 ELISA (DY6220-05, R&D Systems) according to manufacturer’s instructions.

### Immunofluorescence (IF) staining

Fixed organoids were embedded in optimal cutting temperature (OCT) compound, frozen, and sectioned at 5 μm. Thawed slides were blocked with 5% bovine serum albumin (BSA, Sigma-Aldrich) in PBS for 1 hour at room temperature then probed with anti-fibroblasts antibody, clone TE-7 (1:200 dilution, NBP2-50082, Bio-Techne, Minneapolis, USA) and anti-AQP5 (1:100 dilution, A9927, Abclonal, Woburn, USA) diluted in 2.5% BSA in PBS overnight at 4°C. Slides were washed with PBS with 0.2% Tween-20 (PBS-T, Sigma-Aldrich) three times. Goat anti-rabbit IgG conjugated to Alexa Fluor 488 (1:200 dilution, A-11008, Thermo Fisher Scientific) and goat anti-mouse IgG conjugated to Alexa Fluor 594 (1:200 dilution, A-11005, Thermo Fisher Scientific) was diluted to 2.5% BSA in PBS and incubated for 1 hour at room temperature. Nuclei were stained with 1 μg/mL DAPI in PBS, mounted using ProLong Gold Antifade mountant (Thermo Fisher Scientific), and imaged using the EVOS M5000 microscope (Thermo Fisher Scientific).

### Bulk RNA-sequencing

Ribosomal RNA depletion RNA sequencing was performed by Canada’s Michael Smith Genome Sciences Centre, Vancouver, Canada, according to an established standard operating procedure. Libraries were sequenced on an Illumina NovaSeq X Plus using paired-end 150-base reads, targeting 100 million reads per library. Library preparation and sequencing procedures are detailed in the Supplementary Methods.

### RNA-seq Quantification and Analysis

Reads were processed using the nf-core/rnaseq pipeline (version 3.26.0) executed with Nextflow (version 26.04.3), following the nf-core/rnaseq usage documentation [31,32]. The STAR-Salmon route was used, in which reads were aligned to the genome with STAR (version 2.7.11b) and transcript-level abundances were estimated with Salmon (version 1.10.3) against the GENCODE v49 GRCh38 primary assembly and annotation [33].

Salmon estimates were imported into R with tximport and summarized to gene-level counts [34]. Counts were filtered using filterByExpr, normalized by trimmed mean of M-values method, and analyzed using limma-voom with the design formula batch + condition. Contrasts included TGF-β1 versus control, bleomycin versus control and TGF-β1 versus bleomycin. Batch-adjusted expression values were used only for visualization. Statistical analyses were performed using the unadjusted counts with batch included as a model covariate. Differential expression was defined by a Benjamini-Hochberg adjusted P value below 0.05 and an absolute log2 fold change above 1. Gene-set enrichment analysis was performed with fgsea using genes ranked by the limma moderated t-statistic. Hallmark gene sets were obtained from MSigDB using msigdbr, and Benjamini–Hochberg correction was applied within each collection and contrast. All results were generated in reproducible, version-tracked R sessions (R version 4.5.0). R package versions are given in the supplementary material. Analysis code is available at https://github.com/COERDUBC/Tri-culture-alveolar-organoid.git. Authors used Claude Code (Anthropic) to organize and validate the analysis code, and the authors reviewed and verified all output and take full responsibility for the content of this publication.

### Targeted transcriptional analysis

A prespecified basal module (Supplementary Table 3) was derived from the aberrant-basaloid IPF-severity program reported by Ryu et al. (2024) and was tested against the moderated-t ranking of each of the contrasts [19]. Normalized enrichment scores, nominal P-values, and leading-edge genes were reported. A targeted marker panel adapted from Li et al. (2026) [35] was also examined. Marker plots show batch-adjusted voom log2 expression and genome-wide adjusted P values from the corresponding limma-voom contrasts.

### CIBERSORTx analysis

A five-state single-cell reference containing AT1, AT2, aberrant basaloid, fibroblast and myofibroblast cells was constructed from the control, chronic obstructive pulmonary disease (COPD), and IPF donors in GSE136831 [16]. The reference was used to generate a CIBERSORTx signature matrix [36]. CIBERSORTx was run in absolute mode without quantile normalization and with S-mode correction. The resulting reference-derived abundance scores were analyzed using the same batch + condition design, with Benjamini-Hochberg correction applied within each contrast. Reference sampling procedures and cell counts are provided in the Supplementary Methods.

### Ligand-receptor analysis

Candidate ligand–receptor pairs were obtained from ConnectomeDB2025 [37] and restricted to pairs when both genes expressed at least 1 count per million or greater in at least half of the organoid libraries. Sender and receiver states were assigned using the GSE136831 single-cell reference, and only pairs assigned to different cell states were retained. NicheNet was used to rank ligands according to their recovery of the contrast-specific differential-expression response. The 20 highest-ranked ligands from each contrast were retained for visualization. Chord plots display the 40 pairs with the highest reference-expression scores, calculated as the geometric mean of ligand and receptor expression in their assigned sender and receiver states.

## Results

### Co-culture of three cell types in an organoid environment was viable using optimized co-culture media

To generate tri-culture alveolar-like organoids, NCI-H441, MRC-5, and CI-huArlo cells needed to be maintained within a shared culture environment. Since these three cell types are typically expanded in different media formulations based on manufacturer recommendations and our own optimization, we first assessed whether terminal-airway organoid medium (TOM) could support the viability of each cell type. A resazurin-based viability assay was performed after 24 and 48 hours of incubation in different media formulations. Across all three cell types, TOM supported viability at levels comparable to, or higher than, their respective expansion media, supporting its use for subsequent tri-culture organoid experiments (**Supp. Fig. 2**).

**Figure 2.**
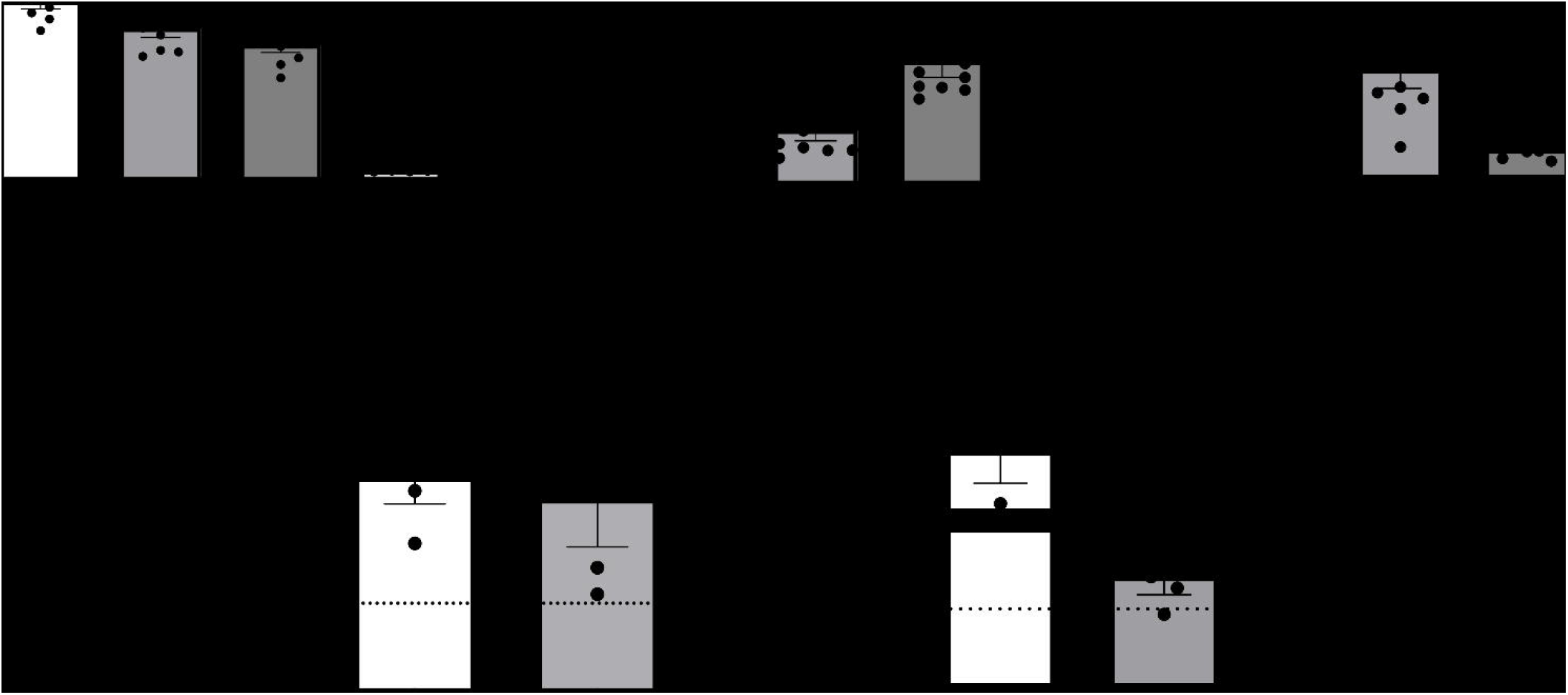
TGF-β1 and bleomycin induce distinct fibrosis-associated phenotypes in tri-culture lung organoids. Organoids were treated with TGF-β1 at 50 ng/mL or bleomycin at 20 µg/mL. **(A)** Metabolic activity as a measure of cellular viability was assessed using resazurin fluorescence across vehicle control, TGF-β1, and bleomycin-treated organoids. The negative control was included with DMSO-treated organoids. **(B)** IL-8 release into the culture supernatant was assayed using ELISA. **(C)** Pro-collagen I release was assayed using ELISA following TGF-β1 or bleomycin-treatment. **(D and E)** Relative *CDKN1A* and *COL1A1* gene expression was assessed by qPCR and calculated using the 2−ΔΔCq method relative to untreated controls. Individual data points represent independent replicates presented as mean ± standard error of the mean. The dotted line in D and E indicates the expression level of the untreated control. Ns, not significant, **P < 0.01, ***P < 0.001

Following this optimization, the three cell lines were expanded in their respective culture medium, mixed at a ratio of 11:7:4 of NCI-H441:CI-huArlo:MRC-5 cells and resuspended in 90% GelTrex Flex diluted with TOM. The cell suspension was dispensed as 30 μL domes. Following dome formation, the organoids were cultured in TOM supplemented with 2.5% GelTrex Flex. Organoid growth was monitored over time. Representative images show progressive expansion, with representative images (**Figure 1A**) showing gradual expansion of organoids, accompanied by an increase in average diameter (**Figure 1B**) and a decrease in organoid counts in each dome (**Figure 1C**). After 2 weeks in culture, IF staining showed diffuse expression of aquaporin-5 (AQP5), a marker for alveolar epithelial type I cells, and TE-7, a marker specific for fibroblasts, showing fluorescent signal along the periphery of each organoid (**Figure 1D**). Hematoxylin and eosin histological staining showed a hollow organoid with the presence of cell debris in the middle of the organoids (**Figure 1E**). Together, these findings demonstrate that the tri-culture system generated three-dimensional alveolar-like structures containing both epithelial cells and fibroblasts.

### Organoids stimulated with TGF-β1 and bleomycin induce distinct inflammatory and ECM responses

Tri-culture organoids were treated with TGF-β1 at 50 ng/mL or bleomycin at 20 µg/mL for 48 hours as two distinct methods to induce features of pulmonary fibrosis in these organoids. Following treatment, a resazurin-based viability assay was performed to assess for cellular metabolic activity as an indirect estimate of cell viability. Fluorescence intensity measured at 560 nm excitation and 590 nm emission, did not differ significantly between TGF-β1-treated organoids and control or bleomycin-treated organoids, indicating no detectable change in viability. However, bleomycin-treated organoids had significantly lower fluorescence intensity than controls (**Figure 2A**), indicating decreased viability. Supernatants were collected following treatment and were assayed for interleukin-8 (IL-8) release as a measure of pro-inflammatory paracrine signalling, and pro-collagen-I as a functional indicator of active type I collagen synthesis and secretion. IL-8 release was significantly increased in bleomycin-treated organoids compared with both TGF-β1-treated organoids and controls, while no significant difference was observed between the TGF-β1 and control groups (**Figure 2B**). In contrast, pro-collagen I release was significantly increased following TGF-β1 treatment compared with both bleomycin-treated organoids and controls, with no significant difference between the bleomycin and control groups (**Figure 2C**). RNA analysis on organoids revealed that *CDKN1A* expression did not differ significantly between the TGF-β1 and bleomycin treatment groups (**Figure 2D**). However, *COL1A1* expression was approximately 20-fold higher in TGF-β1-treated organoids than in bleomycin-treated organoids (**Figure 2E**).

### TGF-β1 and bleomycin treatment induced distinctive transcriptomic responses in organoids

Bulk RNA-sequencing was performed to characterize the transcriptomic responses of tri-culture alveolar-like organoids. Principal component analysis (PCA) demonstrated separation among control, TGF-β1-treated, and bleomycin-treated organoids with PC1 and PC2 accounting for 29% and 13% of the total transcriptional variance, respectively (**Supp. Figure 3A**). No overlap among the three groups was observed. Hierarchical clustering of sample distances reproduced this grouping and indicated that TGF-β1 and bleomycin induced distinct yet reproducible transcriptional responses (**Supp. Figure 3B**). Relative to controls, TGF-β1-treatment resulted in 750 significantly upregulated and 641 downregulated genes (**Supplementary Table 1**). Genes showing strong upregulation included *COL1A1, COL4A1, COL14A2, FN1, TGFB1, VCAN, MMP2, ADAM12, ADAM19,* and *SERPINE*2, consistent with increased signatures associated with extracellular matrix production, cell adhesion and cytoskeletal remodelling (*ITGAV, ITGB6, TNS1, PALLD*, and *TPM1*) [38–41], and positive autoregulation of TGF-β1 signalling (*TGFB1, LEFTY2, SKIL,* and *DACT1*) [42–46]. (**Figure 3A**). Hallmark gene set enrichment analysis (GSEA) identified positive enrichment of pathways related to epithelial-mesenchymal transition, TGF-β1 signalling, apical junction, angiogenesis, and myogenesis programs. Interferon alpha and gamma responses, cholesterol homeostasis, and fatty acid metabolism showed negative enrichment. Together, these findings show that TGF-β1 treatment produces a transcriptional response dominated by extracellular matrix remodeling and profibrotic signalling.

**Figure 3.**
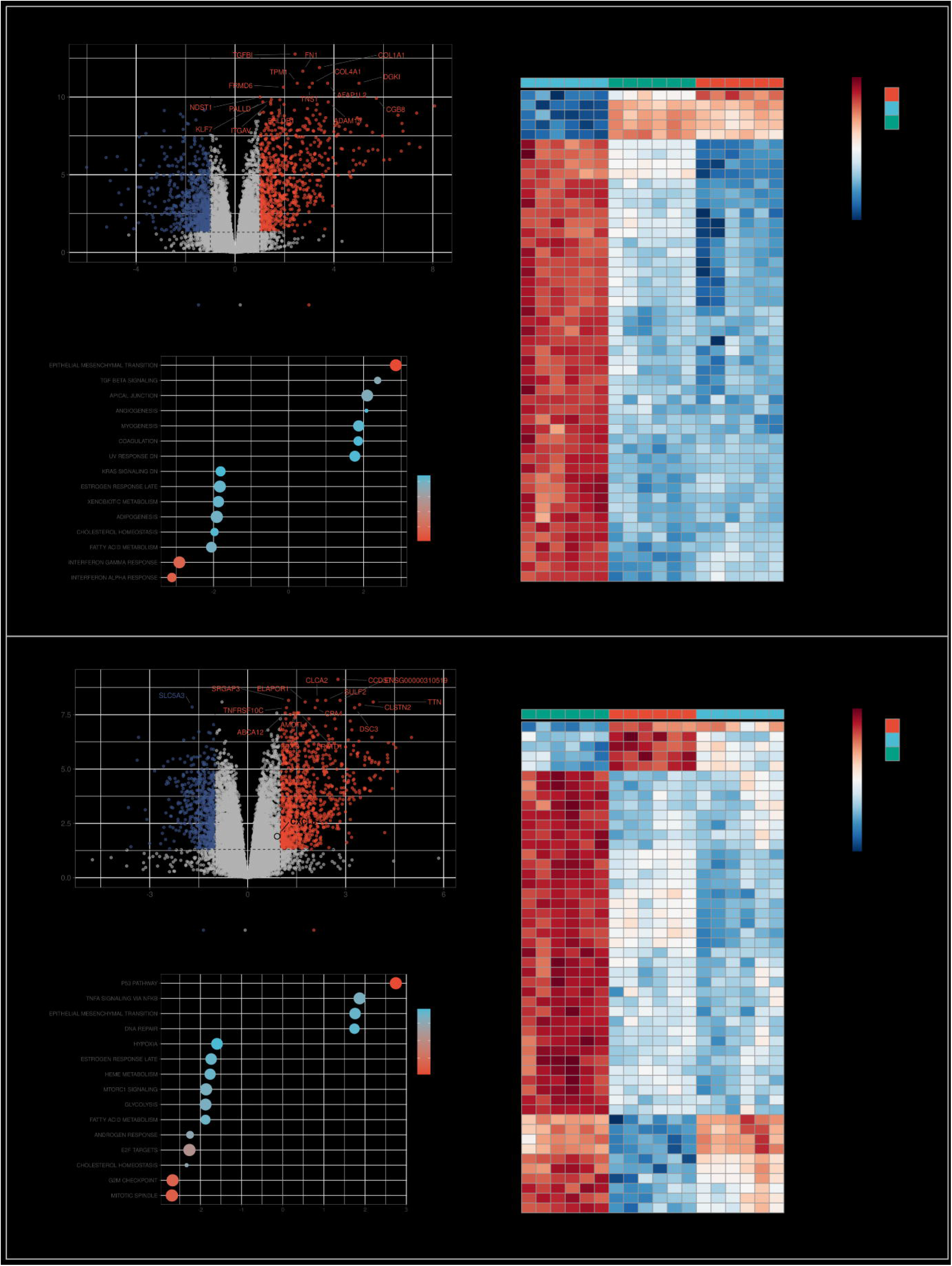
TGF-β1 and bleomycin induce different transcriptional programs in alveolar-like organoids. **A.** Differential gene expression was assessed between TGF-β1-treated **(A)** or bleomycin-treated **(B)** and untreated control organoids per condition using a limma–voom model including batch and condition as covariates (∼ batch + condition). Volcano plots show log2 fold changes and genome-wide BH-adjusted P-values, highlighting genes with FDR < 0.05 and |log2FC| > 1. Heatmaps show row-scaled expression of the top 50 genes from each contrast, while Hallmark GSEA summarizes positively and negatively enriched pathways. Positive NES indicates higher expression in the lefthand condition of contrast, and negative indicates higher expression in the right-hand condition.

Bleomycin treatment resulted in 953 significantly upregulated and 552 significantly downregulated genes relative to untreated controls (Supplementary Table 1). Among the genes most significantly upregulated following bleomycin treatment, several were associated with the DNA damage response, p53 signaling, and cell cycle regulation, including *CDKN1A*, *FDXR*, *CLCA2*, *TNFRSF10C*, *XPC*, and *POLH*. Consistent with these changes, pathway analysis showed positive enrichment of p53 signalling and DNA repair pathways and negative enrichment of E2F target, G2M checkpoint, and mitotic spindle pathways. Bleomycin also increased *DSC3*, *ABCA12*, and *ALOXE3*, genes associated with epithelial adhesion and lipid-barrier differentiation, as well as the lipid associated genes, *PLIN4, ABCA1,* and *PLTP*.

### TGF-β1 and bleomycin treatment drives organoids towards aberrant basaloid and myofibroblast states

To determine whether profibrotic stimulation shifted organoids toward the epithelial and mesenchymal states observed in human fILD, bulk RNA-sequencing profiles were analyzed using CIBERSORTx with reference signatures derived from the human lung IPF Cell Atlas (GSE136831) [16]. Five reference cell states were evaluated, including AT1, AT2, aberrant basaloid, fibroblast, and myofibroblast states (**Figure 4**). Compared with control organoids, TGF β1 treated organoids showed higher aberrant basaloid and myofibroblast scores and lower AT1 and AT2 scores. Their fibroblast scores were also higher than control but lower than those observed following bleomycin treatment. Bleomycin treated organoids showed the highest fibroblast scores and lower AT2 scores than controls, whereas their aberrant basaloid, AT1, and myofibroblast scores remained closer to control values.

**Figure 4.**
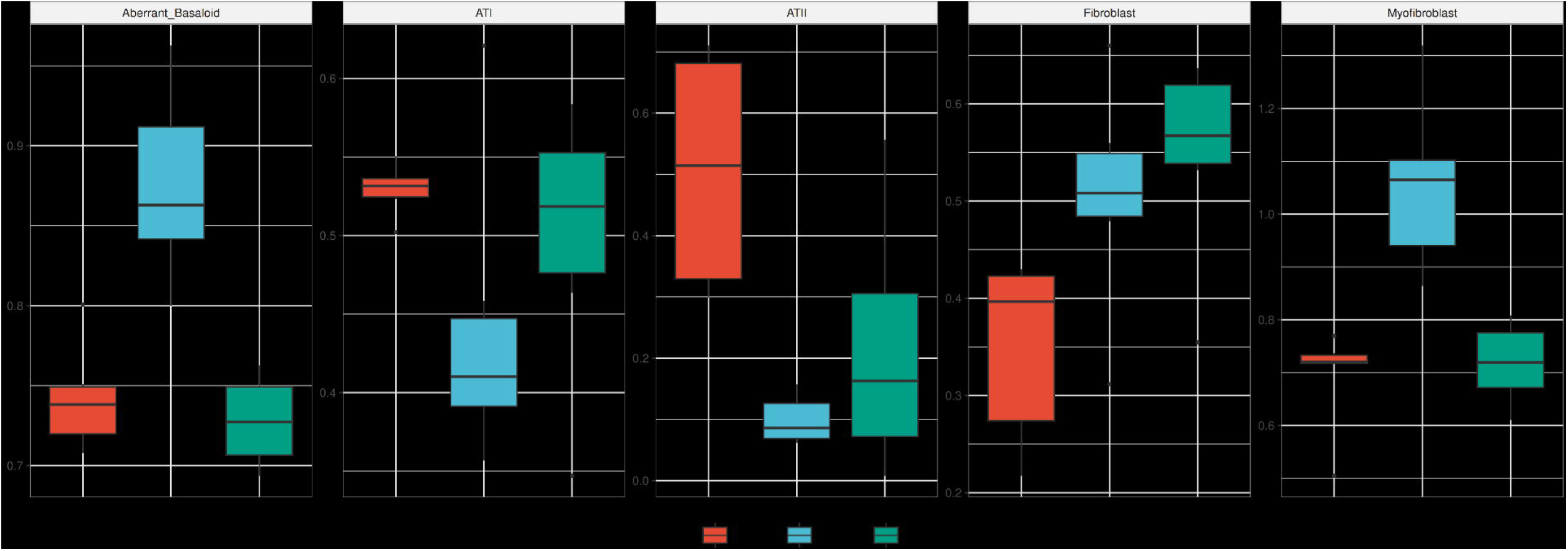
Cellular deconvolution for five expected cell states in the organoids. Absolute CIBERSORTx scores for aberrant basaloid, AT1, AT2, fibroblast, and myofibroblast reference signatures were compared among control, bleomycin, and TGF-β1 treated organoids. Boxes represent the interquartile range, centre lines indicate medians, and whiskers extend to 1.5 times the interquartile range, and points indicate outliers.

### TGF-β1 accounts for most predicted ligand-receptor interactions from mesenchymal senders

Treatment-associated ligand-receptor candidates were mapped for the five reference states. The resulting chord diagram demonstrated extensive predicted connectivity between epithelial and mesenchymal states following treatment (**Figure 5**). Recurrent connections involving S100A4 and TIMP1 suggested that these ligands could participate in several cell-state-specific communication pathways through receptors which include *AGER, ANXA2, CD74, CD63, ITGB1*, and *LRP1*. Among the 30 shortlisted cell-state-specific interactions, 25 were associated only with TGF-β1 treatment, four were shared between TGF-β1 and bleomycin, and one was specific to bleomycin. Mesenchymal states were the predominant predicted senders, accounting for 25 of the 30 interactions, including 17 originating from cells with a myofibroblast signature and 8 from those with a fibroblast signature.

**Figure 5.**
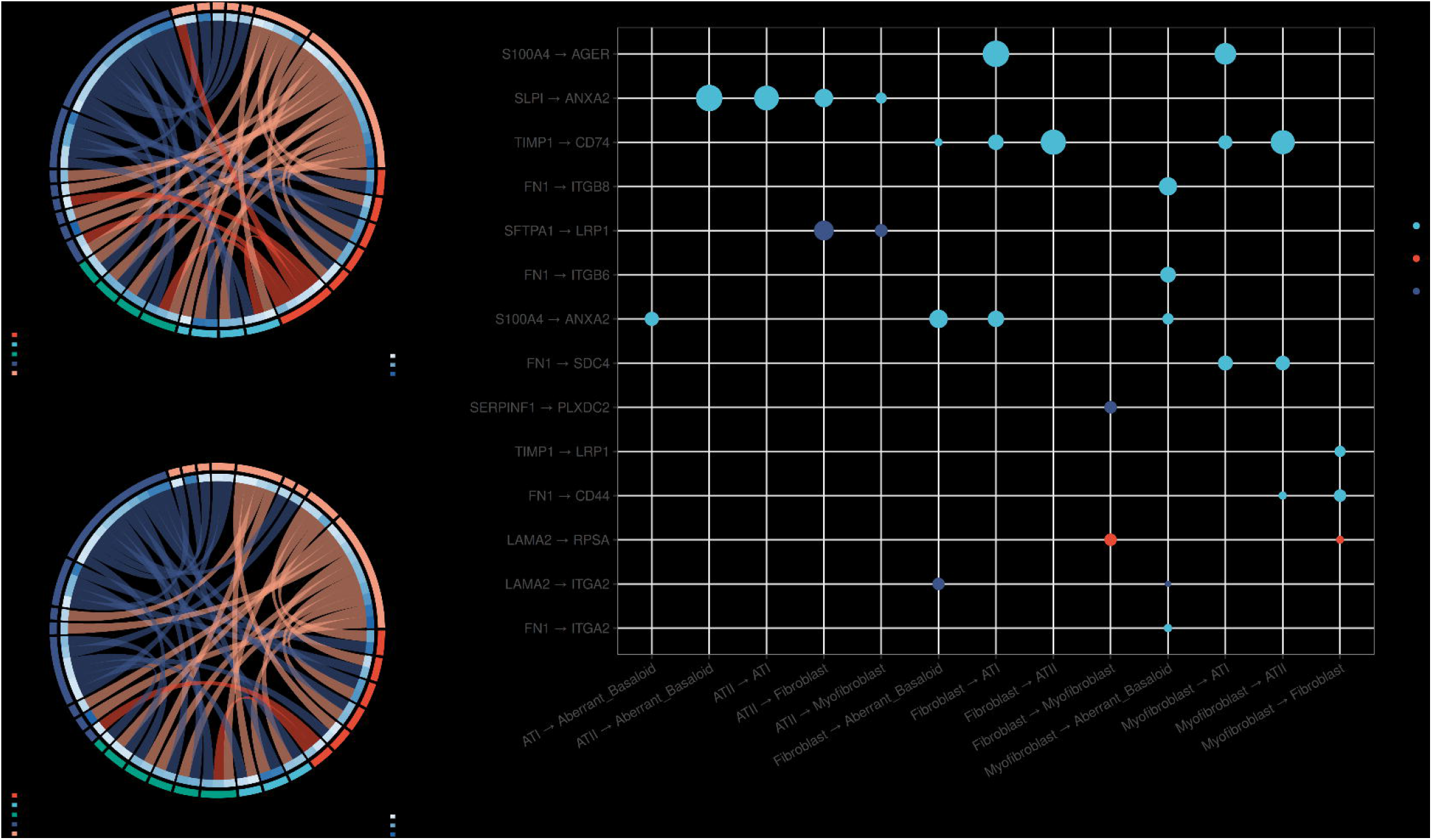
NicheNet-prioritized ligand–receptor candidates mapped to GSE136831 reference epithelial and mesenchymal states. (A,B) Chord diagrams show treatment-associated candidate interactions for TGF-β1 and bleomycin. Ribbon colour denotes the ligand-producing sender state, outer sector colour denotes cell state, and ribbon width and the inner blue ring represent the GSE136831 reference-expression score. **(C)** Bubble plot shows the 30 highest-scoring treatment-associated pairs, with colour indicating treatment association and size indicating reference score. These are computationally prioritized candidates, not experimentally validated interactions

### Comparison with published aberrant-basaloid and alveolar-injury signatures

To determine whether treatment induced a severity-associated basal transcriptional program, a prespecified 18 gene module was evaluated by GSEA. Sixteen genes were retained after expression filtering, while *LGALS7B* and *CALML3* fell below the expression filter (**Figure 6**). The module was enriched in bleomycin treated organoids relative to controls (NES = 1.50, nominal P = 0.048). The leading-edge subset comprised of *DSC3, GJB5, S100A2, LY6D*, and *SERPINB5*, all of which had positive log2 fold changes of 3.27, 1.46, 0.70, 0.71, and 0.71 respectively. The module was similarly shifted toward bleomycin in the TGF-β1 versus bleomycin contrast, although this did not meet the nominal significance threshold (P = 0.084). Eleven of the 16 genes were expressed more highly following bleomycin than TGF-β1 treatment and included the downregulation of *DSC3* (log2FC = −3.70) in TGF-β1.

**Figure 6.**
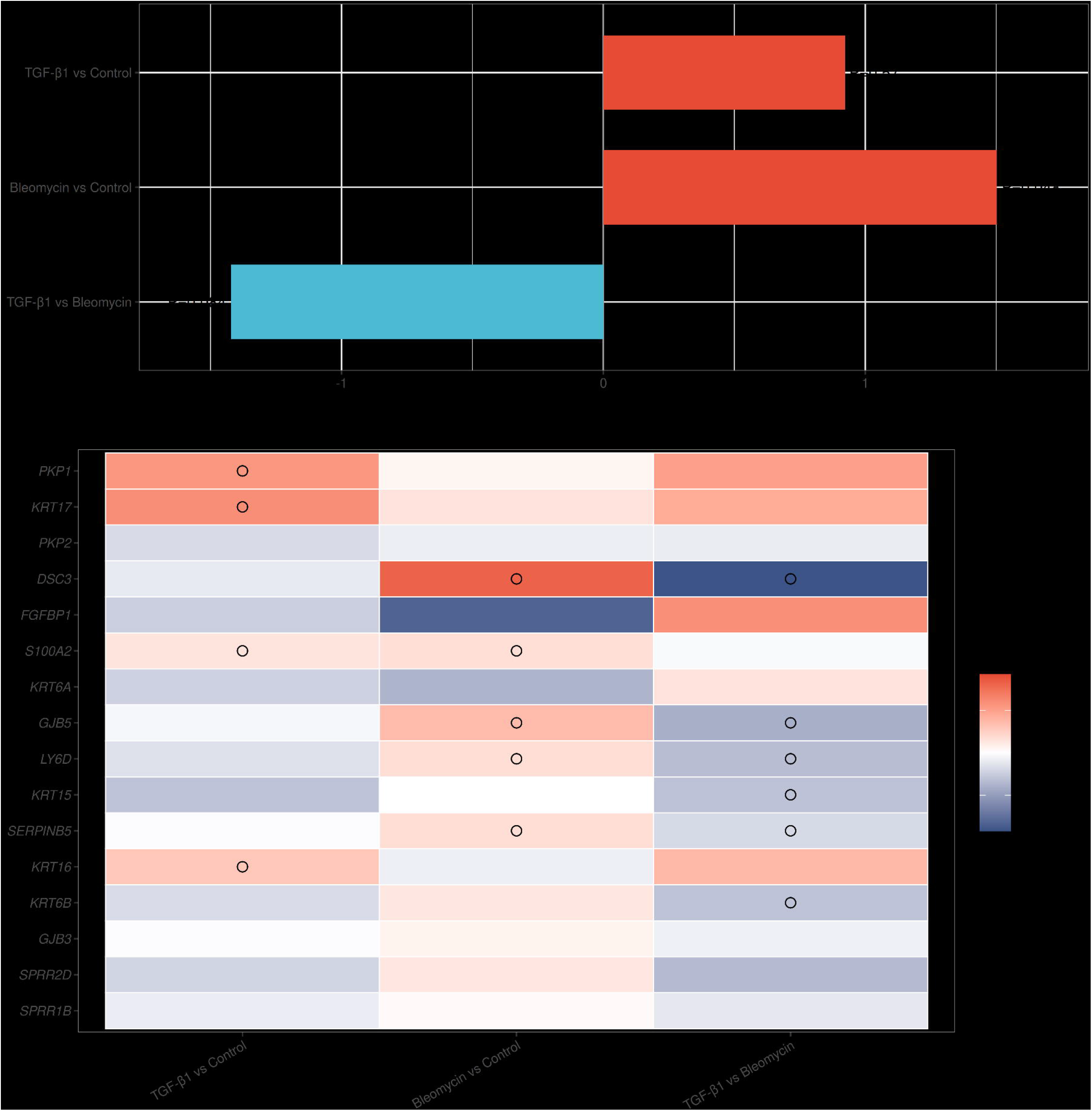
Enrichment of a basal module from the aberrant-basaloid IPF-severity programme. **(A)** Normalized enrichment score (NES) for the module in each contrast, tested by fgsea against limma-voom moderated-t rankings. A positive NES indicates higher expression in the lefthand condition of contrast, and negative indicates higher expression in the right-hand condition. Labels are nominal P values. **(B)** limma log2 fold change (log2FC) for all 18 module genes (rows) in each contrast (columns). Positive log2FC indicates higher expression in the lefthand condition of contrast, and negative indicates higher expression in the right-hand condition. Circles marks leading-edge genes. *LGALS7B* and *CALML3* did not pass the expression filter.

TGF-β1 treatment did not produce coordinated enrichment of the module compared to control, though four upregulated genes *PKP1, KRT16, KRT17,* and *S100A2* (log2FC = 2.21, 1.19, 2.39, 0.59) were observed.

A curated marker panel adapted from Li et al. was then used to compare the tri-culture response with primary human AT2-derived spheroids treated with TGF-β1, the hypoxia mimic dimethyloxalylglycine (DMOG), or both (**Figure 7**) [35]. Because the studies differed in cellular composition, treatment conditions, readouts, and duration, this comparison was limited to qualitative marker-level concordance. *SFTPC* was excluded because it fell below the expression filter. Among the seven markers evaluated under TGF-β1 treatment, five showed concordant responses: *KRT17, COL1A1*, and *CDH2* were upregulated (log2FC = 2.39, 3.42, and 1.27, respectively; all FDR < 4 × 10□□), *GDF15* decreased (log2FC = −0.91, FDR = 0.0015), and *CDKN1A* remained unchanged. *MMP7* and *AGER* were unchanged, and HES4 was unchanged under both treatments. TGF-β1 therefore reproduced a partial aberrant epithelial-marker response, without the *AGER* induction or AT1-like transition. Although there was no DMOG treatment included in this study, *SLC2A1* increased modestly after TGF-β1 treatment (log2FC = 0.43, FDR = 4.65 × 10□□), aligned with the positively enriched Hallmark hypoxia pathway (NES = 1.62, FDR = 0.0023). Bleomycin increased *CDKN1A* and *GDF15* (log2FC = 1.18 and 1.54, respectively; all FDR < 4 × 10□□), and modestly reduced MMP7 (log2FC = −0.76, FDR = 0.019), whereas *KRT17, COL1A1, CDH2,* and *AGER* were unchanged. This profile supported a stress and growth arrest response containing senescence-associated markers, although it did not establish cellular senescence. Together, the module and marker analyses distinguished a coordinated basal injury program with senescence-associated signalling following bleomycin treatment from a selective TGF-β1 response involving *KRT17* and matrix and adhesion associated genes.

**Figure 7.**
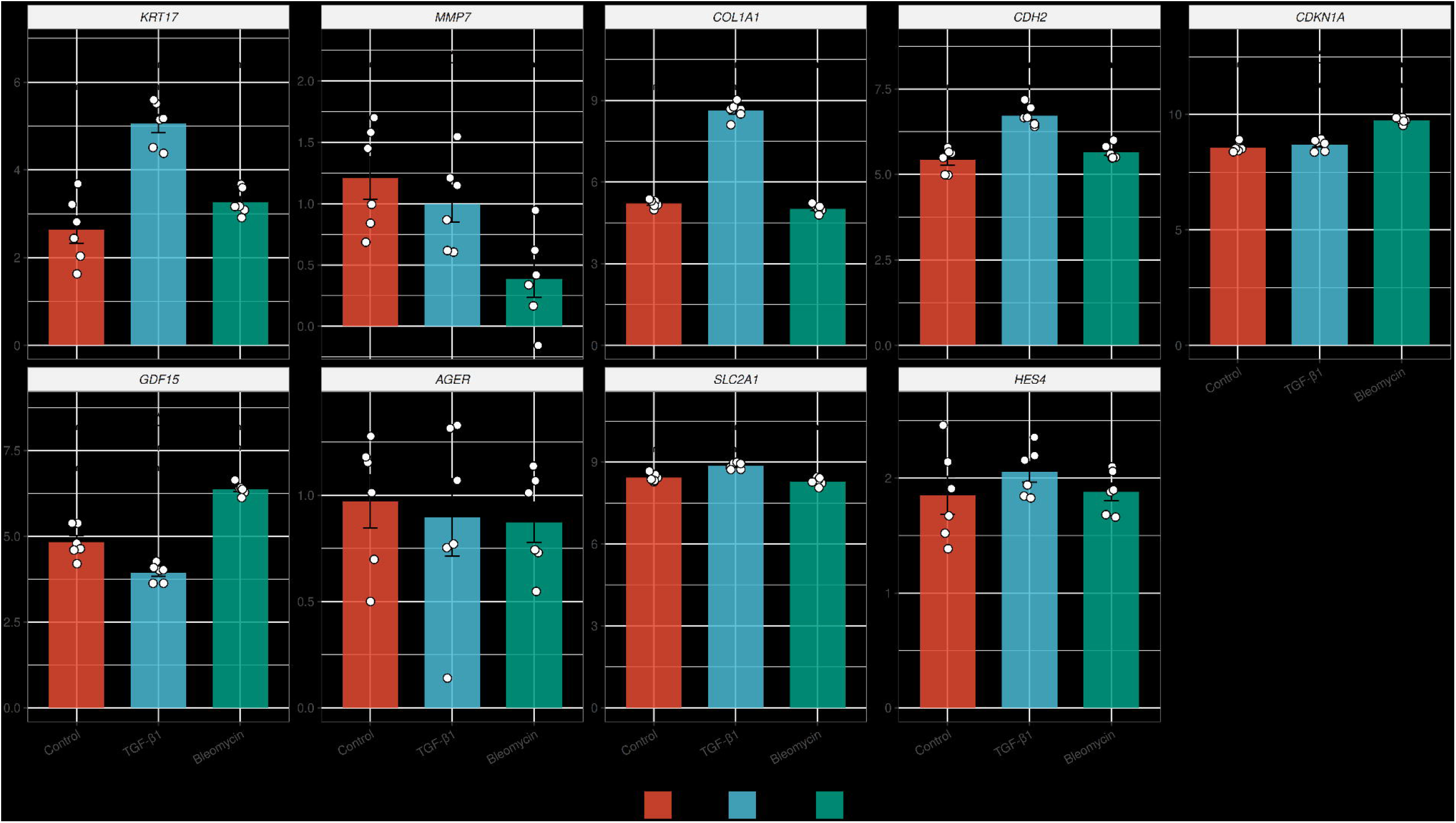
Fibrosis and alveolar marker expression. Batch-adjusted voom log2 expression of a nine-gene panel. Gene list was adapted from Li et al. (2026) [35]. Bars are means ± SEM, open circles individual libraries (n = 6 per condition), y-axes are panel specific. Brackets use genome-wide Benjamini–Hochberg-adjusted limma-voom P values (*** = p<0.001; ** = p<0.01; * = p<0.05). *SFTPC* fell below the expression filter and is omitted.

## Discussion

This study developed and characterized a three-dimensional alveolar-like organoid model composed of NCI-H441 and CI-huArlo epithelial cells with MRC-5 fibroblasts. The three cell types formed multicellular structures containing AQP5-positive epithelial and TE-7-positive fibroblast populations. TGF-β1 and bleomycin produced distinct responses across metabolic, secreted-protein, targeted gene-expression, and transcriptomic measurements. TGF-β1 induced a matrix-dominant response characterized by increased COL1A1 expression, procollagen I secretion, and extracellular matrix-related transcriptional programs. Bleomycin reduced metabolic activity, increased IL-8 secretion, and induced DNA-damage, inflammatory, and epithelial-remodelling programs without increasing the measured collagen endpoints. Reference-based analyses identified transcriptional similarities to selected epithelial and mesenchymal states reported in human IPF.

CI-huArlo cells were derived as a single-cell clone of the polyclonal cell line, CI-hAELVi. Previous studies combining CI-hAELVi cells with NCI-H441 cells in air-liquid interface cultures demonstrated epithelial barrier formation and retention of selected surfactant-associated features, including pro-surfactant protein C, ABCA3 expression, and lamellar bodies [27,28,47]. Building on these studies and previous epithelial-mesenchymal work [24,25,28,47,48], we combined NCI-H441, CI-huArlo, and MRC-5 cells at an 11:7:4 ratio to create an epithelial-predominant tri-culture.

In this study, two profibrotic stimuli were used. TGF-β1 contributes to direct profibrotic signalling for fibroblast activation and extracellular matrix production [49]. Bleomycin is a chemotherapeutic agent that induces oxidative stress and DNA strand damage [50]. Compared to TGF-β1, bleomycin can injure the epithelial compartment which contributes to the release of inflammatory and profibrotic signals to activate fibroblasts and is commonly used as an experimental fibrosis-inducer with various *in vitro* and *in vivo* models [51,52]. Following organoid formation, TGF-β1 and bleomycin produced distinguishable phenotypes with TGF-β1 increasing *COL1A1* expression and procollagen I secretion and inducing transcriptional programs associated with extracellular matrix production, TGF-β signalling, adhesion, and contractile remodelling. Bleomycin reduced resazurin reduction, increased IL-8 secretion, and induced transcriptional programs associated with p53, DNA repair, inflammatory, and epithelial remodelling programs while suppressing cell-cycle programs. The separation of control, TGF-β1, and bleomycin-treated groups through PCA and hierarchical clustering further showed that each condition produced a reproducible transcriptomic state.

This model occupies a complementary experimental position relative to primary and pluripotent stem cell-derived alveolar organoids. These systems have been used to examine epithelial injury, fibroblast activation, altered alveolar differentiation, impaired regenerative capacity, and disease-associated genetic phenotypes [51,53–57]. However, these systems usually require long differentiation procedures and more specialized culture conditions. The present model uses established cell sources and a 14-day formation period, which may facilitate experimental scaling and modular addition of defined stimuli or cell populations, and, by combining epithelial and mesenchymal cells, can enable the study of multicellular interactions.

The contrast between TGF-β1 and bleomycin treatment was a central finding. TGF-β1 increased COL1A1 expression and procollagen I secretion, together with expression of COL4A1, FN1, and TGFBI. Enrichment of TGF-β signalling, epithelial-mesenchymal transition, and myogenesis gene sets was consistent with coordinated matrix remodelling, altered cell-matrix interactions, and contractile transcriptional activity [58–60]. These findings agree with the established role of TGF-β1 in fibroblast activation and extracellular matrix production [49]. However, exogenous active TGF-β1 directly engages a major fibrogenic pathway and therefore does not reproduce the upstream epithelial injury, inflammatory signalling, or latent TGF-β activation that can initiate fibrosis in vivo [14,61].

Bleomycin produced a different response. Reduced resazurin reduction and increased IL-8 secretion, indicated cellular injury and inflammatory signalling, while the absence of increased COL1A1 expression or procollagen I secretion suggested that the 48-hour exposure did not generate a comparable matrix response. Positive enrichment of p53 and DNA-repair pathways, increased expression of *CDKN1A* and other p53-responsive genes, and suppression of E2F, G2M-checkpoint, and mitotic-spindle programs were consistent with genotoxic stress and reduced cell-cycle progression [62–64]. Bleomycin has also been shown to induce p21-associated senescence-like phenotypes in alveolar epithelial cells [65] but senescence was not directly measured here. These findings therefore support an acute injury and growth-arrest response rather than demonstrating cellular senescence or established fibrosis. Longer, repeated, or recovery-phase experiments will be required to determine whether recurrent bleomycin injury produces persistent epithelial dysfunction and subsequent matrix remodelling.

Reference-based deconvolution provided a link between the organoid responses and cell states reported in human IPF. Untreated organoids had the highest AT1 and AT2 reference-state scores. Bleomycin reduced both alveolar epithelial scores and produced the highest fibroblast score, whereas TGF-β1 produced the highest myofibroblast and aberrant basaloid scores. These directions are broadly consistent with the loss of normal alveolar epithelial states and emergence of aberrant epithelial and mesenchymal states reported in IPF lungs [16,66,67]. Human pluripotent stem cell-derived alveolar organoids can also be driven toward KRT17-positive transitional states by fibrosis-associated cytokines, and recent fibroblast-dependent organoid studies have used similar states as therapeutic readouts [66,67]. However, CIBERSORTx absolute scores are reference-dependent measures of transcriptional resemblance rather than cell fractions or evidence of lineage conversion [36]. These findings are most appropriately interpreted as associations with enrichment of these cell states rather than direct measurement of abundance or lineage.

The ligand-receptor analysis generated a complementary set of hypotheses about epithelial-mesenchymal communication. Mesenchymal states were the predicted senders in 25 of the 30 shortlisted interactions, including 17 interactions from the myofibroblast reference state and eight from the fibroblast state. Twenty-five interactions were detected only following TGF-β1 exposure, four were shared between TGF-β1 and bleomycin, and one was specific to bleomycin. This distribution was consistent with the stronger matrix and myofibroblast-like response to TGF-β1. The most directly coherent network involved ECM ligands, including FN1 and COL1A1, paired with integrins, CD44, and SDC4, which could coordinate cell adhesion and matrix sensing. Recurrent non-matrix candidates included S100A4-AGER, TIMP1-CD74, and epithelial SLPI-ANXA2 interactions. Physical interactions between S100A4 and RAGE, TIMP1 and CD74, and SLPI and annexin A2 have been demonstrated in other cellular systems and disease contexts [68–70]. However, their roles in alveolar epithelial-fibroblast communication and lung fibrosis remain unknown, and further investigation is required.

The module and marker analyses examined related but distinct features of injury associated epithelial remodelling. The module test assessed enrichment of a severity-associated basal program derived from IPF lung tissue, whereas the marker panel compared individual injury-responses with those reported in primary AT2-derived spheroids. KRT17 is the only shared gene between these two tests. Bleomycin produced two complementary transcriptional signals, enrichment in the severity-associated basal module and induction of stress and growth arrest markers *CDKN1A* and *GDF15*. However, TGF-β1 treatment increased *COL1A1, CDH2* and *KRT17* at the marker level but left the module unchanged, suggesting concurrent activation of basal-associated and senescence-associated programs rather than a single, basaloid program [71], since TGF-β1 needs BMP inhibitor to promote transdifferentiation in AT-2 cells [57]. Bleomycin engaged DNA damage and epithelial senescence, features carried by aberrant basaloid cells in the IPF lung and the inferred abundance of this population is the cell type most strongly associated with worsened FVC and DLCO in IPF [19]. Together these results indicate that the two stimuli model different parts of the disease: TGF-β1 reproduces the matrix response with only a partial basal shift, whereas bleomycin reproduces the senescence-associated programme that tracks IPF severity. Longer or repeated exposures and direct senescence readouts would be needed to establish whether a basaloid-like state is genuinely formed.

This study has two principal limitations. First, the cellular composition provides a reproducible but simplified representation of the fibrotic alveolar niche. The organoids were generated from adenocarcinoma-derived NCI-H441 cells, immortalized CI-huArlo cells, and fetal lung MRC-5 fibroblasts. These sources do not reproduce the aging, genetics, telomere dysfunction, epigenetic alterations, and pre-existing cellular abnormalities found in patients with IPF [72]. The model here also lacks immune and endothelial populations that can regulate fibroblast proliferation, myofibroblast activation, extracellular matrix turnover, and angiocrine signalling during fibrosis [73,74]. Although this reductionist approach limits reconstruction of the complete fibrotic alveolar niche, they permit more controlled investigation of epithelial-fibroblast responses, with future experiments able to optimize organoid and tissue-engineered models that incorporate these additional populations [20].

Second, the experimental approaches support an acute, stimulus-specific response rather than the establishment of chronic fibrosis. Exogenous active TGF-β1 directly activates a major downstream fibrogenic pathway, bypassing endogenous cytokine production, latent TGF-β1 activation, and upstream injury-dependent mechanisms involved in fibrosis [61,76]. Furthermore, bleomycin acts as an injury-inducing and DNA-damaging agent where under the 48 hour exposure used here, bleomycin reduced metabolic activity and increased IL-8 release without increasing *COL1A1* expression or secreted procollagen I, suggesting a predominantly acute cellular injury and inflammatory signalling state than fibrogenesis, and treatment was limited to a 48-hour exposure at a single concentration, which is congruent with regimens used by other groups [77–79], but leads to an injury and matrix response indicative of an acute, stimulus-specific response rather than chronic progressive fibrosis. Other features that future experiments should be done to further characterize the response to examine deposited and mature ECM, such as assessing matrix stiffness and fibrillar collagen deposition, myofibroblast differentiation associated with increase alpha smooth muscle actin expression as well as cellular and gel contraction and presence of persistent or progressive fibrosis independent of continued treatment through longer culturing post-treatment and a wash-out.

Several features strengthen the experimental platform. The study incorporated two epithelial cell types and one fibroblast type within the same three-dimensional matrix, used a shared medium that supported each cellular compartment, and assessed responses across morphology, metabolic activity, secreted proteins, targeted gene expression, and transcriptome-wide profiling. The use of six biological replicates per RNA-sequencing condition and the concordance between clustering, differential expression, pathway enrichment, and biochemical measurements increase confidence that TGF-β1 and bleomycin generated distinct responses. The model is also modular. It could be used to compare acute and repeated exposures, incorporate mechanical strain, add immune or endothelial populations, and test pathway inhibitors within a controlled background.

This study establishes a defined tri-culture alveolar-like organoid platform that distinguishes a TGF-β1-associated matrix response from a bleomycin-associated injury response. Its transcriptional resemblance to selected human IPF-associated states provides testable hypotheses rather than evidence that the model reproduces IPF pathology. With cell-resolved validation, direct assessment of deposited matrix and alveolar function, and comparison with primary donor-derived systems, the platform may support mechanistic studies of epithelial-mesenchymal interactions in fibrotic lung injury.

## Conclusions

This tri-culture alveolar-like organoid model distinguishes a TGF-β1-associated matrix response from a bleomycin-associated acute injury response, providing a defined platform for studying epithelial-fibroblast interactions during fibrotic lung injury.

## Declarations

### Ethics approval and consent to participate

N/A

## Consent for publication

N/A

## Availability of data and materials

Analysis code is available at https://github.com/COERDUBC/Tri-culture-alveolar-organoid.git. The repository is private during peer review and access will be provided to the editors and reviewers on request. It will be made publicly available and archived with a Zenodo DOI upon publication.

## Authors’ contributions

Conceptualization: T.J.F.G., I.T.S.L., M.H.R., and E.T.O.

Methodology, validation, and formal analysis: T.J.F.G. and Y.M.

Software: Y.M. and M.H.R.

Investigation: T.J.F.G., Y.M., S.K., T.M., A.M., K.S.K.L., C.G., M.R.P., and C.P.S.L.

Data curation: T.J.F.G., S.K., T.M., A.M., K.S.K.L., C.G., M.R.P., and C.P.S.L.

Writing, original draft: T.J.F.G. and Y.M.

Writing, review and editing: T.J.F.G., Y.M., S.K., T.M., A.M., K.S.K.L., C.G., M.R.P., C.P.S.L., G.C.G., A.J.H., C.J.R., C.C., M.H.R., E.T.O.

Visualization: T.J.F.G.

Supervision: G.C.G., A.J.H., C.J.R., C.C., I.T.S.L., M.H.R., and E.T.O.

Funding acquisition: M.H.R. and E.T.O.

## Acknowledgements

We would like to thank the members of the Osei and Ryu laboratories for their thoughtful contributions. The authors wish to acknowledge Canada’s Michael Smith Genome Sciences Centre, Vancouver, Canada for Ribosomal RNA depletion RNA sequencing.

---

**Supplemental Figure 1. Representative image of CI-huArlo, NCI-H441, and MRC-5 cells embedded in a GelTrex dome.** 13,200 total cells were used at a ratio of 11:7:4 between NCI-H441, CI-huArlo, and MRC-5 cells with each dome required 3.5 μL of TOM and 31.5 μL of GelTrex. These domes were dispensed into each well of the prewarmed plate and solidified at 37°C.

**Supplemental Figure 2. Terminal-airway Organoid Medium (TOM) supports the metabolic activity and viability of the three constituent cell lines.** NCI-H441, MRC-5, and CI-huArlo were incubated for 24 or 48 hours in DMEM or RPMI-1640, both supplemented with 10% FBS, PneumaCult-Ex, HuAEC medium, TOM, or PBS as a negative control. Cellular metabolic activity as an indirect measurement for cellular viability was assessed using a resazurin-based assay. Bars represent mean ± standard error of the mean, with each individual data point representing a biological replicate. One-way ANOVA followed by Tukey’s multiple-comparisons test was performed with significance annotations representing * = p<0.05, ** = p<0.01, *** = p<0.001, and **** = p<0.0001.

**Supplemental Figure 3. TGF-β1 and bleomycin induce distinct global transcriptional responses in tri culture alveolar like organoids.** Bulk RNA sequencing was performed on untreated control organoids and organoids treated for 48 hours with bleomycin at 20 μg/mL or TGF-β1 at 50 ng/mL. **A.** Principal component analysis of batch corrected expression data. PC1 and PC2 accounted for 29% and 13% of the total transcriptional variance, respectively. Each point represents one RNA sequencing library. **B.** Heatmap of pairwise sample distances with hierarchical clustering. Lower distances are represented by darker blue. Samples clustered according to treatment condition, with six biological replicates included per condition.

**Supplemental Table 1.** Differentially expressed genes in each treatment versus Control.

**Supplemental Table 2.** Full list of the significantly differentially expressed genes in each treatment versus Control.

**Supplemental Table 3.** The 18 prespecified basal module genes from Ryu et al. 2024.

**Supplemental Table 4.** Cell sampling summary for the GSE136831 single-cell reference.

