## Supplementary Materials for "Characterization of tri-culture alveolar-like organoids to study fibrotic lung diseases"

**Supplemental Methods**

**Organoid harvest, RNA extraction, optimal cutting temperature (OCT) processing, and viability assay**

After 48 hours of incubation, the supernatant was collected, centrifuged at 500 x g for 5 minutes at 4°C to clear it of debris, aliquoted, and stored at -80°C until analysis. The organoid domes were washed with warmed PBS, then 350 μL of 10% PrestoBlue (Thermo Fisher Scientific), a resazurin-based dye, in TOM was added. Domes were incubated for 2 hours at 37°C, and supernatants were transferred onto a standard 96-well plate and read on a spectrophotometer plate reader (Molecular Devices) with excitation at 560 nm and emission at 590 nm. The domes were washed with warmed PBS; organoid harvesting solution (R&D Systems) was added to break up the domes and placed on ice. The tubes were finger vortexed every 5 minutes until the organoids were floating in the solution then pelleted at 250 x g for 5 minutes at 4°C. The supernatant was discarded, and the resulting pellet was washed and resuspended with cold PBS before pelleting with the same conditions again. Organoid pellets were either lysed with Monarch StabiLyse DNA/RNA buffer (New England Biolabs, Ipswich, USA) and stored at -80°C temporarily until processing with the Monarch Total RNA Miniprep kit (New England Biolabs) or lysed with working lysis buffer from the MagMAX mirVana Total RNA isolation kit (Thermo Fisher Scientific) and processed using the KingFisher Apex purification system (Thermo Fisher Scientific). For OCT embedding, the organoid dome was fixed with 1% paraformaldehyde in PBS for 20 minutes at room temperature before being washed with PBS. 20 μL of the organoid dome was also aspirated and dispensed gently in a small droplet of OCT compound (Fisher Scientific, Waltham, USA) dispensed onto a disposable base mold (VWR, Radnor, USA) and flash frozen in a dry-ice bath containing isopentane (Sigma-Aldrich). The mold was then filled with OCT compound until full and flash frozen again and stored at -80°C until cryo-sectioning. Sections were cut using a cryostat (Leica Biosystems, Nussloch, Germany) at 5 μm and stored at -80°C.

**Quantitative PCR and immunoassays**

Lysed organoid pellets were thawed on ice and total RNA was isolated using the Monarch Spin RNA isolation kit (New England Biolabs) according to manufacturer’s instructions. Complementary DNA was synthesized from 20 to 600 ng of RNA using the High-Capacity cDNA Reverse Transcription Kit according to manufacturer’s instructions (Thermo Fisher Scientific). Quantitative PCR was performed on a QuantStudio 6 Pro Real-Time PCR system (Thermo Fisher Scientific) using relative quantification in fast mode. Each 10 μL reaction contained TaqMan Fast Advanced Master Mix, TaqMan Gene Expression Assays, and 300 ng of cDNA. GAPDH-VIC (Assay ID: Hs99999905_m1) was used as the endogenous reference assay, with forward (5′-CAAGGTCATCCATGACAACTTTG-3′), and reverse (5′-GGCCATCCACAGTCTTCTGG-3′) primers as previously reported [24]. COL1A1-FAM (Assay ID: Hs00164004_m1) and CDKN1A-FAM (Assay ID: Hs00355782_m1) were used as target assays. The thermal cycling conditions consisted of an initial hold at 95°C for 20 seconds, followed by 40 cycles of 95°C for 1 second and 60°C for 20 seconds. Relative gene expression was calculated by normalizing target gene expression to GAPDH.

**Bulk RNA-sequencing**
To remove cytoplasmic rRNA and mitochondrial rRNA transcripts from total RNA, NEBNext rRNA Depletion Kit for Human/Mouse/Rat was used (New England Biolabs, E7405X). Enzymatic reactions were set-up in a 96-well plate (Thermo Fisher Scientific) on a Microlab NIMBUS liquid handler (Hamilton Robotics, Nevada, USA). 100-500 ng of DNase I-treated total RNA in 6 µL was hybridized to rRNA probes in an 8.5 µL reaction. Heat-sealed plates were incubated at 95°C for 2 minutes followed by incremental reduction in temperature by 0.1°C per second to 22°C (730 cycles). The rRNA in DNA hybrids were digested using RNase H in an 11 µL reaction incubated in a thermocycler at 50°C for 30 minutes. To remove excess rRNA probes (DNA) and residual genomic DNA contamination, DNase I was added in a total reaction volume of 26 µL and incubated at 37°C for 30 minutes. RNA was purified using RNA MagClean DX beads (Aline Biosciences, Woburn, USA) with 15 minutes of binding time, 7 minutes clearing on a magnet followed by two 70% ethanol washes, 5 minutes to air dry the RNA pellet and elution in 18µL DEPC water. The plate containing RNA was stored at -80°C prior to cDNA synthesis. First-strand cDNA was synthesized from the purified RNA (minus rRNA) using the Maxima H Minus First Strand cDNA Synthesis kit (Thermo Fisher, USA) and random hexamer primers at a concentration of 8 ng/µL along with a final concentration of 0.04 µg/µL Actinomycin D, followed by PCR Clean DX bead purification on a Microlab NIMBUS robot (Hamilton Robotics). The second strand cDNA was synthesized following the NEBNext Ultra Directional Second Strand cDNA Synthesis protocol (NEB) that incorporates dUTP in the dNTP mix, allowing the second strand to be digested using USERTM enzyme (NEB) in the post-adapter ligation PCR and thus achieving strand specificity. cDNA was fragmented by Covaris LE220 sonication for 130 seconds (2x65 seconds) at a “Duty cycle” of 30%, 450 Peak Incident Power (W) and 200 Cycles per Burst in a 96-well microTUBE Plate (P/N: 520078) to achieve 200-250 bp average fragment lengths. The paired-end sequencing library was prepared following Canada's Michael Smith Genome Sciences Centre strand-specific, plate-based library construction protocol on a Microlab NIMBUS robot (Hamilton Robotics) using the NEBNext Ultra II DNA Library Prep Kit (New England Biolabs). Briefly, the sheared cDNA was subject to end-repair and dA-tailing in a single reaction using the End Prep Enzyme mix (NEB), incubated at 20oC for 30 minutes. Ligation to TruSeq adapters was performed at 20oC for 15 minutes. The adapter-ligated products were purified using PCR Clean DX beads followed by indexed PCR using NEBNext Ultra II Q5 Master Mix (New England Biolabs) with USER enzyme (1 U/µL, NEB) and dual indexed primer set. PCR parameters: 37˚C for 15 minutes, 98˚C for 1 minute followed by 13 cycles of 98˚C for 15 seconds, 65˚C for 30 seconds and 72˚C for 30 seconds, and then 72˚C for 5 minutes. The PCR products were purified and size-selected using a 0.9:1 PCR Clean DX beads-to-sample ratio, and the eluted DNA quality was assessed with Agilent DNA 1000 assay (Agilent Technologies, Santa Clara, USA) or Caliper LabChip GX for DNA samples using the High Sensitivity Assay (PerkinElmer, Shelton, USA) and quantified using a Quant-iT dsDNA High Sensitivity Assay Kit on a Qubit fluorometer (Invitrogen) prior to library pooling and size-corrected final molar concentration calculation for Illumina NovaSeq X Plus sequencing with paired-end 150 base reads on a 25B flowcell targeting 100 million reads per library.

**Supplemental Figures**

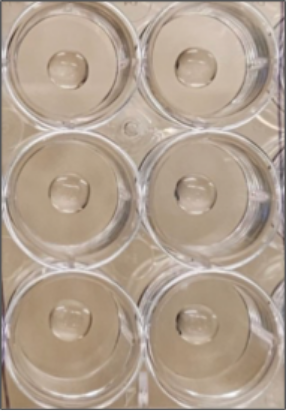

**Supplemental Figure 1. Representative image of CI-HuARLO, NCI-H441, and MRC-5 cells embedded in a GelTrex dome.** 13,200 total cells were used at a ratio of 11:7:4 between NCI-H441, CI-huArlo, and MRC-5 cells with each dome required 3.5 μL of TOM and 31.5 μL of GelTrex. These domes were dispensed into each well of the prewarmed plate and solidified at 37°C.

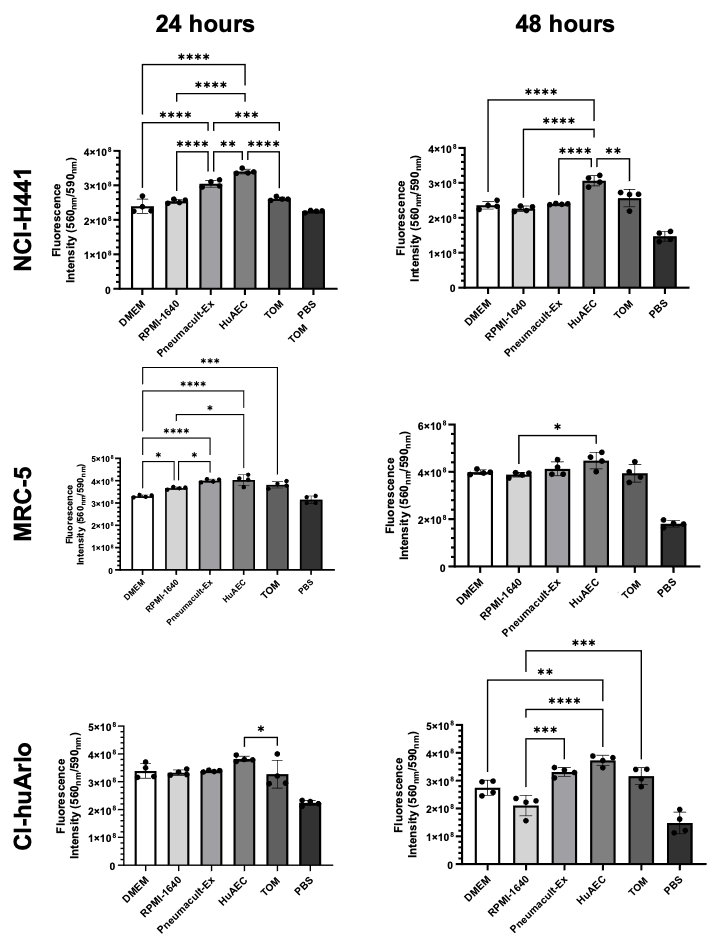

**Supplemental Figure 2. Terminal-airway Organoid Medium (TOM) supports the metabolic activity and viability of the three constituent cell lines.** NCI-H441, MRC-5, and CI-HuARLO were incubated for 24 or 48 hours in DMEM or RPMI-1640, both supplemented with 10% FBS, PneumaCult-Ex, HuAEC medium, TOM, or PBS as a negative control. Cellular metabolic activity as an indirect measurement for cellular viability was assessed using a resazurin-based assay. Bars represent mean ± standard error of the mean, with each individual data point representing a biological replicate. One-way ANOVA followed by Tukey’s multiple-comparisons test was performed with significance annotations representing * = p<0.05, **= p<0.01, *** = p<0.001, and **** = p<0.0001.

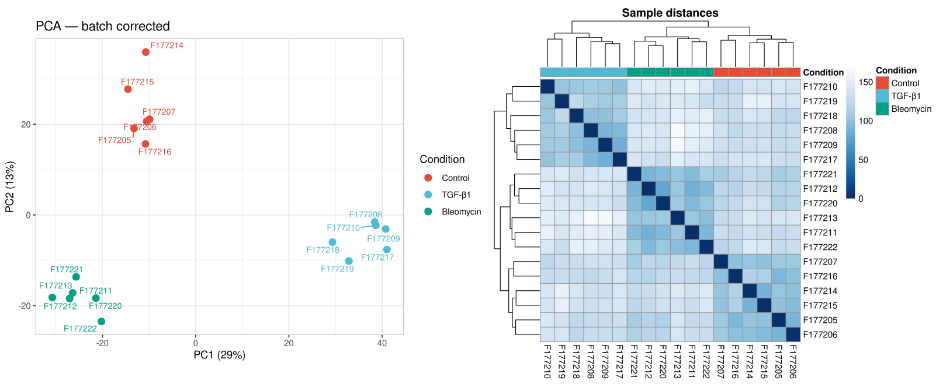

**Supplemental Figure 3. TGF-β1 and bleomycin induce distinct global transcriptional responses in tri culture alveolar like organoids.** Bulk RNA sequencing was performed on untreated control organoids and organoids treated for 48 hours with bleomycin at 20 μg/mL or TGF-β1 at 50 ng/mL. **A.** Principal component analysis of batch corrected expression data. PC1 and PC2 accounted for 29% and 13% of the total transcriptional variance, respectively. Each point represents one RNA sequencing library. **B.** Heatmap of pairwise sample distances with hierarchical clustering. Lower distances are represented by darker blue. Samples clustered according to treatment condition, with six biological replicates included per condition.

**Supplemental Table 1.** Differentially expressed genes in each treatment versus Control.

**Supplemental Table 2.** Full list of the significantly differentially expressed genes in each treatment versus Control.

**Supplemental Table 3.** The 18 prespecified basal module genes from Ryu et al. 2024.

**Supplemental Table 4.** Cell sampling summary for the GSE136831 single-cell reference.
